# A “feet digging” swash zone sampling method for the sandy beach bivalve *Donax deltoides* (Bivalvia: Donacidae)

**DOI:** 10.1101/686196

**Authors:** Stephen Totterman

## Abstract

A “feet digging” method for sampling the sandy beach bivalve *Donax deltoides* was evaluated by comparison to quadrat-based results from eleven beaches in subtropical eastern Australia. The method was developed from a recreational fishing technique that involves twisting one’s feet into the thixotropic sand to dislodge buried clams which are then recovered by hand. Several plots are sampled across the swash zone in one five-minute sampling unit and the process is replicated at several locations along the beach. Mean feet digging clam counts were proportional to mean transect linear clam densities (*r* = 0.98). Clam length-frequency distributions from feet digging were similar to those from quadrat sampling except that feet digging was not effective for clams < 16 mm. Feet digging counts are sensitive to both across shore (tidal) and alongshore variation in clam abundance and were less precise than those from quadrat-based methods (*CV* 1.2× larger). However, feet digging is fast and the method should be useful for low cost surveys of *Donax deltoides* and similar “surf clams”.

## 1. Introduction

This study compares sampling methods for *Donax deltoides*. Commonly known as the “pipi”, *D. deltoides* is a large surf clam found along the eastern and southern coasts of Australia, from Fraser Island, Queensland, across New South Wales (NSW) and Victoria, to the Murray River, in South Australia. *D. deltoides* grows to a maximum length of 75 mm and can develop large populations on some beaches, where they are harvested by commercial and recreational fishers (Ferguson *et al.* 2014). Owing to its large size, *D. deltoides* can make up a large component of the macrobenthic biomass over a wide range of densities (*e.g.* McLachlan *et al.* 1996) and has an important role in the ecology of sandy beaches. Efforts to quantify *D. deltoides* populations have recently increased following growth in commercial and recreational fisheries for this species (Gray *et al.* 2014; Ferguson *et al.* 2015).

Fishery-based catch per unit effort (CPUE) is generally a poor index of abundance for species like *D. deltoides* that form dense aggregations because local (“patch scale”) catch rates can remain high despite large decreases in the population (beach scale) abundance, *i.e.* “hyperstability” of CPUE (Maunder *et al.* 2006). A further problem is that fishery-based CPUE can only be estimated where catch records are available.

Research sampling of surf clams commonly involves excavating and sieving quadrats or cores on across shore transects at low tide. Integration of quadrat densities (clams/m^2^) across the transect length (m) gives along shore linear densities (clams/m; Totterman 2019a). The single integrated density estimate per transect emphasises that the sampling unit is transect and that quadrats are pseudoreplicates (Millar and Anderson 2004). Linear density estimates absolute abundance and can be used to estimate absolute population size (*i.e.* mean linear density from a sample of transects × beach length).

For fixed area transect designs, areal density (clams/m^2^) or mean counts (clams/transect) can be reported. Areal density should be regarded as a relative abundance estimator because it does not account for beach width and the across shore distribution of levels. Totterman (2019a) demonstrated with simulations that areal density is approximately inversely proportional to transect length. Areal density should not be used to estimate absolute population size. A further problem is that relative abundance results from different surveys and studies that employ different sampling methods and designs are generally not comparable.

Surf clams commonly aggregate in discontinuous “belts” or “bands” with unimodal across-shore distributions (*e.g.* Totterman 2019b). Clam band sampling commonly uses a fixed number of quadrats and estimates areal (relative) density (*e.g.* Owner and Rohweder 2003). Even though clam band sampling involves relatively few quadrats compared to transects, it has been infrequently used because preliminary sampling is often required to locate the band. Recreational fishers occasionally search for buried *D. deltoides* in the intertidal zone at low tide by dragging a knife blade through the sand. James and Fairweather (1995) showed this knife method was effective for locating *D. deltoides* bands with densities > 0.5 clams per 0.1 m^2^. However, the knife method was unsatisfactory for estimating relative abundance because it was sensitive to buried pebbles and insensitive to clams < 30 mm.

Quadrat-based sampling is laborious and this often leads to insufficient replication of transects (Murray-Jones 1999). Three rapid sampling methods have previously been developed for *D. deltoides*. James and Fairweather (1995) proposed a “finger raking” method for recovering clams from flooded quadrats that involves both tactile detection and dislodgment of buried clams, which tend to float to the surface. Finger-raking recovered 98% of clams in comparison to excavation and sieving of quadrats and was rapid except for fetching buckets of water. Finger raking, as applied by James and Fairweather (1995), is simply a modification for quadrat-based sampling.

Gray *et al.* (2014) proposed a 30-second swash zone “hand digging” method where sand and clams are scooped into a 19 mm square mesh net bag. Ferguson *et al.* (2015) used a “cockle rake” with a 44 mm mesh to sample 3 × 1.5 m quadrats in the swash zone. These latter two swash zone sampling methods are CPUE methods, where hand digging effort is time-based and cockle raking effort is area-based. For CPUE data it is usually assumed that catch is proportional to true abundance (Maunder *et al.* 2006). Gray *et al.* (2014) and Ferguson *et al.* (2015) did not compare their CPUE results to quadrat-based estimates of abundance.

This study evaluated a swash zone “feet digging” method for sampling *D. deltoides* in comparison to transect and clam band sampling.

## 2. Materials and Methods

### 2.1 Study sites

Eleven beaches were sampled in northern NSW from the Tweed River to the Wooli River (Table 1; Figure 1). These were the same beaches that were sampled in Totterman (2019b). Beaches where *D. deltoides* was known to be abundant were preferred, including those identified in a previous regional study by Owner and Rohweder (2003). South Ballina, which had the highest and increasing clam abundance was sampled four times (Table 1). Surveys were performed in all months except for March and May. *D. deltoides* was commercially fished on South Ballina and Ten Mile beaches during this study (Gray 2016b, 2016c; pers. obs.).

**Table 1.**
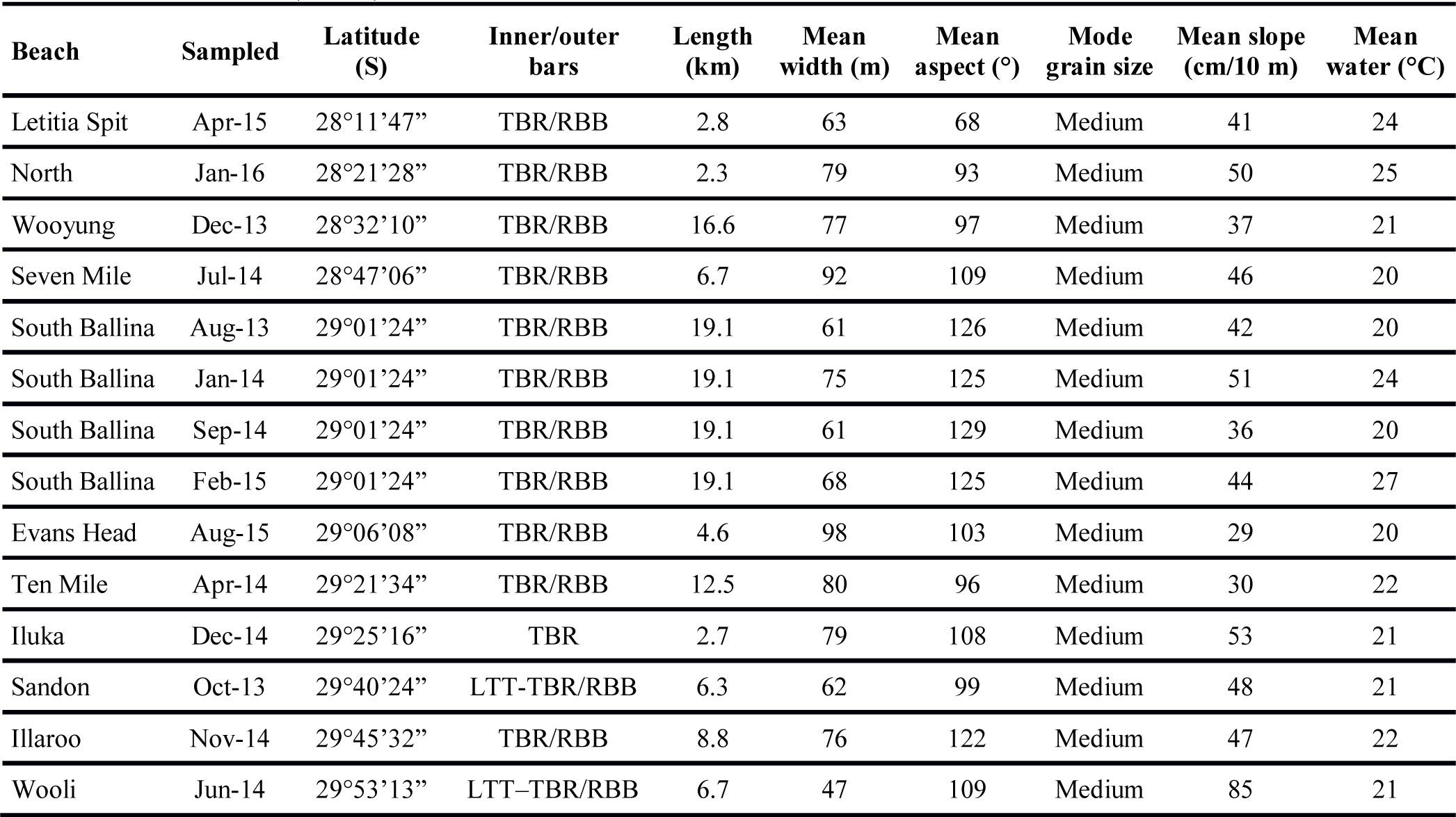
Beach summaries, ordered north to south (Figure 1). Each beach was sampled with 10 transects in the months indicated (South Ballina was sampled on four occasions). Latitude is given for the southern end of each beach. Beach morphodynamics from Short (2007) are, in order of increasing energy: LTT = low tide terrace, TBR = transverse bar and rip, RBB = rhythmic bar and beach. Medium sand has a grain size 0.25–0.5 mm on the Wentworth scale. Measurement of these physical variables is described in Totterman (2019b).

**Figure 1.**
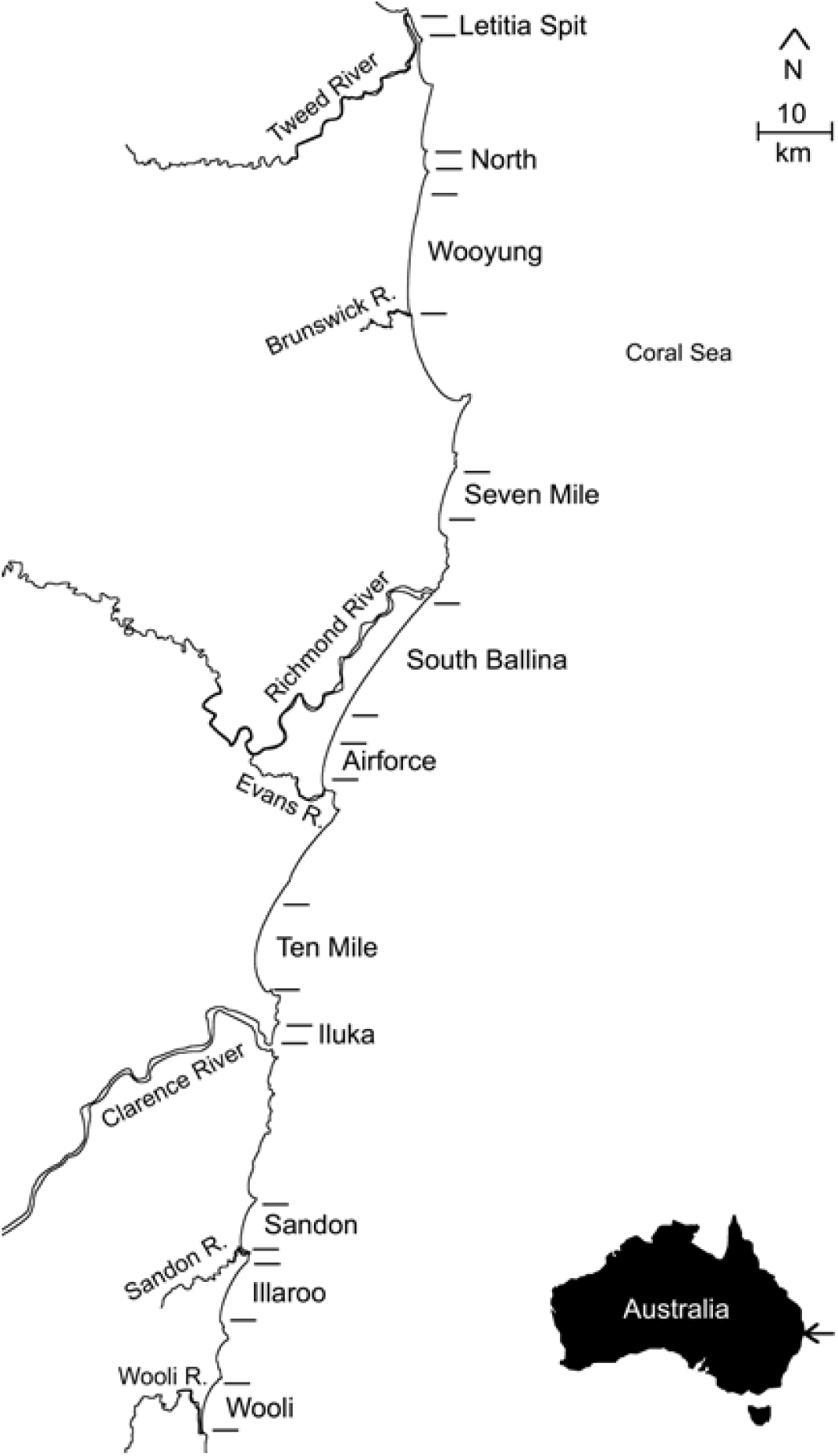
Location of 11 study beaches. The inset map shows the study region on the east coast of Australia (arrowed).

### 2.2 Transect sampling

Each beach was sampled with ten transects, equally spaced along the length of the beach starting at a random offset from the southern end (distance 0 km). These were the same transects that were sampled in Totterman (2019b). Sampling occurred over several days because no more than two transects could be sampled during one low tide period. No particular attention was given towards spring-neap or semi-diurnal variation in the tidal range. *D. deltoides* undergo tidal migrations and seem to optimise their across shore location relative to varying beach conditions (Totterman 2019b).

A fixed area transect design with variable across shore level intervals was used to consistently position transect levels relative to the limits of the intertidal and swash zones. Transects extended from the most recent high tide drift line to the bottom of the low tide swash zone (at the average limit of the backwash). Level one was at the drift line, levels 2–4 were equally spaced in the intertidal zone, level five was at the top of the low tide swash zone (at the average limit of swash run up), level six was in the middle of the swash zone and level seven was at the bottom of the swash zone (average limit of backwash) (Figure 2a). Three replicate quadrats were sampled in a direction parallel to the shoreline at each of levels 1–6, with five paces between quadrats. Excavation of large quadrats underwater is impossible and three feet digging plots were sampled in place of quadrats at level seven (see below). Transects were measured with a marked rope.

**Figure 2.**
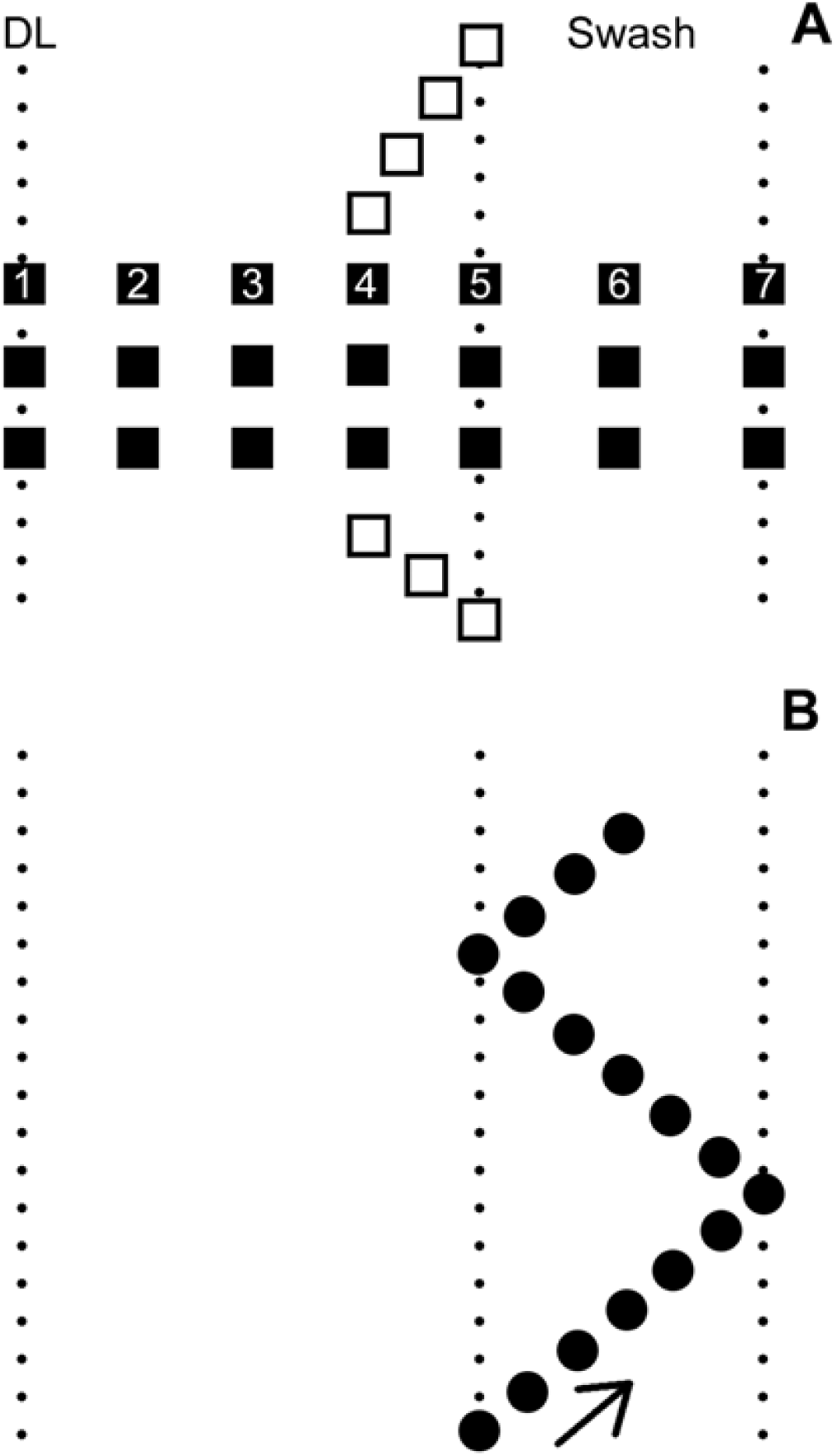
Plan views of transect (A) and feet digging (B) sampling layouts (not to scale). Black squares are transect quadrats. White squares are clam band quadrats (the clam band in this illustration was intersected at transect levels 4 and 5). Black circles are feet digging plots (the arrow indicates the direction of feet digging in this illustration). From left to right, the dotted lines represent the drift line (DL), top of the swash zone and bottom of the swash zone.

A square, sheet metal, 0.1 m^2^ quadrat (sides 31.6 cm long and 10 cm deep) was used (James and Fairweather 1995). Sand was excavated first to 10 cm and then separately from 10–20 cm. A hand shovel was used for gauging the 10–20 cm depth. Excavated material was washed through 6 × 6 millimetre plastic sieves using buckets of seawater except that finger raking (James and Fairweather 1995) was used to rapidly inspect the 10–20 cm depth in saturated sand, where the sides of the quadrat hole were prone to collapsing. Finger raking was also frequently used for the 0–10 cm depth in the swash zone, where waves disrupts sampling efforts. Clams were measured with callipers to the nearest millimetre (maximum shell length) and then released. Clams broken at the edges of the quadrat and not measurable or any clams washed away in the swash zone and lost were added to the count of those measured.

Clam band sampling was performed together with each transect. Quadrats were sampled across the same levels as any clam aggregation intersected by the transect (Figure 2a). If no aggregation was found then clam band sampling was performed at the same levels as any bands present in adjacent transects (Owner and Rohweder 2003). Four replicate quadrats were sampled immediately north of the transect, along an approximately 45 degree diagonal across the middle of the clam band and three quadrats were sampled immediately south (total 4 + 3 = 7 quadrats per band),with five paces between quadrats (Figure 2a).

### 2.3 Feet digging sampling

Feet digging or “feet twisting” is commonly used by recreational fishers collecting *D. deltoides* in the swash zone (see Jaramillo *et al.* 1994 for a previous application of this method). Twisting one’s legs and feet from the hips down while pivoting on the balls of the feet, causes the feet to dig into the sand. Erosion of sand from around the feet by swash and backwash speeds the process. Clams disturbed in the vicinity of the feet rise to the surface where they can be picked up (similar to the finger raking method of James and Fairweather 1995). Large and more firmly anchored clams can be felt with the feet and recovered by hand. The area sampled by one feet digging plot is approximately 0.1 m^2^ and the depth is approximately 10 cm.

Eighty feet digging locations were sampled on each beach, with equal allocation to each of four tidal stages: high (± 2 h), ebb (from high + 2 h to low −2 h), low (± 2 h) and, flow (from low + 2 h to high −2 h). Locations were selected randomly with replacement from sampling grids with 100 m (for short beaches < 4 km) or 200 m along shore intervals. Any repeated locations were sampled on different days.

Feet digging effort was fixed at five minutes per location. Individual feet digging plots were sampled in an approximately 45 degree “zig-zag” pattern parallel to the shore line: 1) dig to approximately ankle depth; 2) move three paces along the diagonal and repeat feet digging; 3) when the bottom/top of the swash zone is reached, change direction across shore and continue feet digging; 4) stop after five minutes (Figure 2b). Five minutes was considered sufficient to sample the width of the swash zone, long enough for timing errors to be negligible and short enough to prevent observer fatigue. Clams were measured and counted as for quadrats. Measuring several hundred or more clams over the course of a sample is time consuming and was statistically unnecessary for this study. On the final sampling occasion for South Ballina, clams were measured only for the first 40 randomly selected feet digging locations, with equal allocation to each of the four tidal stages.

### 2.4 Statistical analysis

The basic sampling unit for this study was the transect/clam band/feet digging location and mean clam abundances were measured at the beach scale (*i.e.* mean density/count per transect/band/feet digging location).

Linear densities were calculated by linear interpolation and integration of level densities across the length of each transect (Totterman 2019a). Relative abundance estimates are reported as mean counts. Scaling mean transect or clam band counts by the transect/band area to obtain densities (clams/m^2^) is unnecessary for fixed area sampling. Mean clam abundances from the three sampling methods were compared pairwise across beaches using scatterplots, Pearson’s product moment correlations and ordinary least squares regression.

Relative standard error (*i.e. RSE* = standard error of mean ÷ mean) is an appropriate measure of precision when comparing methods with different measurement scales and sample sizes (Andrew and Mapstone 1987). For some target *RSE*, the ratio of estimated sample sizes for two sampling methods is the squared ratio of their coefficients of variation (*CV*):

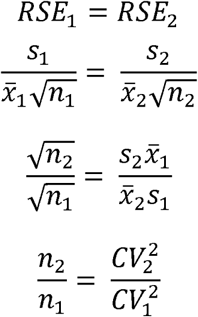

where *n*, 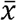 and *s* are the sample size, mean and standard deviation respectively. Sample size ratios among the three sampling methods were computed pairwise within beaches and then mean sample size ratios across all beaches were computed.

For beaches with > 30 clams measured for each sampling method, length-frequency distributions were compared pairwise within beaches using *k*-sample Anderson-Darling tests (Scholz and Stephens 1987). Like the two-sample Kolmogorov-Smirnov test, the *k*-sample Anderson-Darling test compares empirical cumulative distribution functions and is sensitive to differences in shape, location and scale between samples. *P*-values from these multiple comparisons were adjusted using the procedure of Holm (1979).

All statistical analyses were performed using R version 3.3.2 (R Core Team 2016). Anderson-Darling tests used the R package kSamples version 1.2.4 (Scholz and Zhu 2016).

## 3. Results

Correlations between transect, clam band and feet digging mean counts and linear densities were linear, strong (*r* ≥ 0.96) and with zero intercepts (*P* ≥ 0.06) (Figure 3). Excluding the higher density and influential South Ballina data, these correlations remained linear, strong (*r* ≥ 0.80) and with zero intercepts (*P* ≥ 0.22).

**Figure 3.**
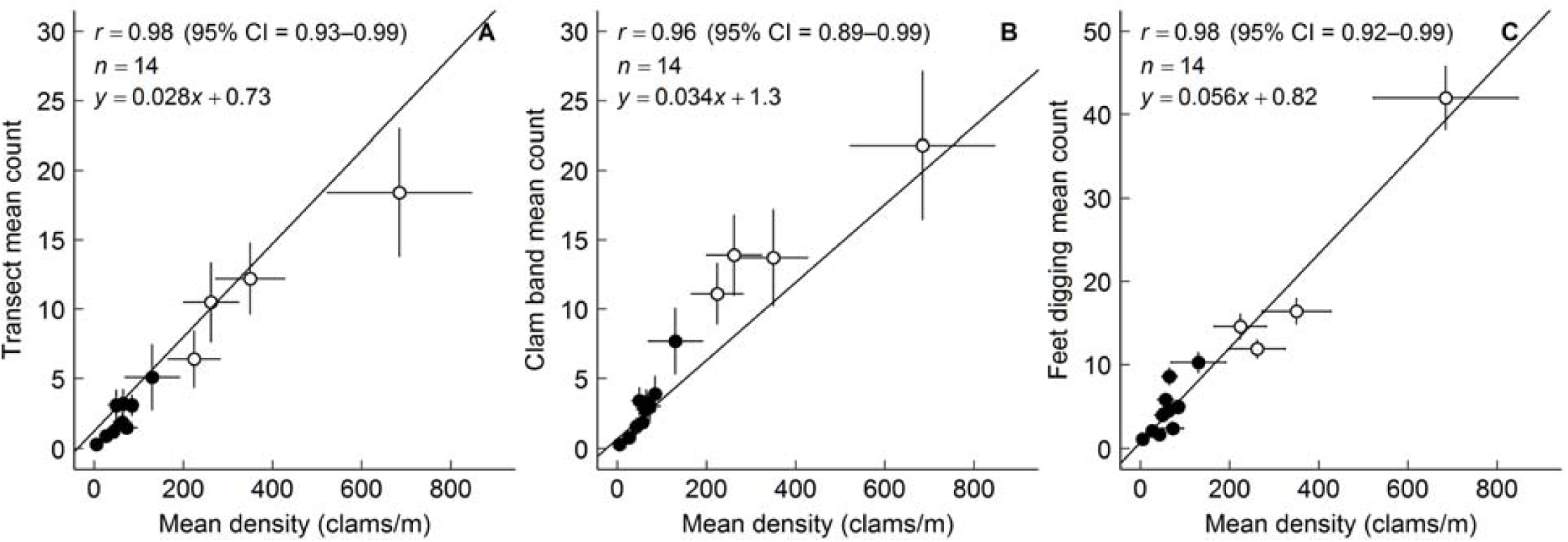
Correlations between three *D. deltoides* relative abundance estimates and transect linear densities. The four South Ballina Beach results (Table 1) are plotted as white circles. Error bars are standard errors. Lines are from ordinary least squares regression.

The along shore distribution of clams was strongly aggregated on all beaches. Variance-mean relationships were approximately quadratic and, with fixed sample sizes, standard errors increased with abundance (Figure 3). Feet digging coefficients of variation averaged 1.2× larger and estimated sample sizes for constant relative standard error averaged 1.6× larger than those for transect linear densities and clam band mean counts.

Feet digging counts indicated that *D. deltoides* was present in the swash zone throughout the tidal cycle (Figure 4). Despite these tidal migrations, seasonal zonation patterns were apparent on some beaches with higher counts occurring towards low tide in summer at South Ballina (Figures 4a, 4b) and Wooyung (Figure 4j) or towards high tide in winter and early spring at Wooli (Figure 4d) and South Ballina (Figures 4f, 4g), but not at Seven Mile (Figure 4e). Correlations between separate high, ebb, low and flow tide feet digging mean counts and linear densities were *r* = 0.82, 0.90, 0.92 and 0.96 respectively (mean *r* = 0.90).

**Figure 4.**
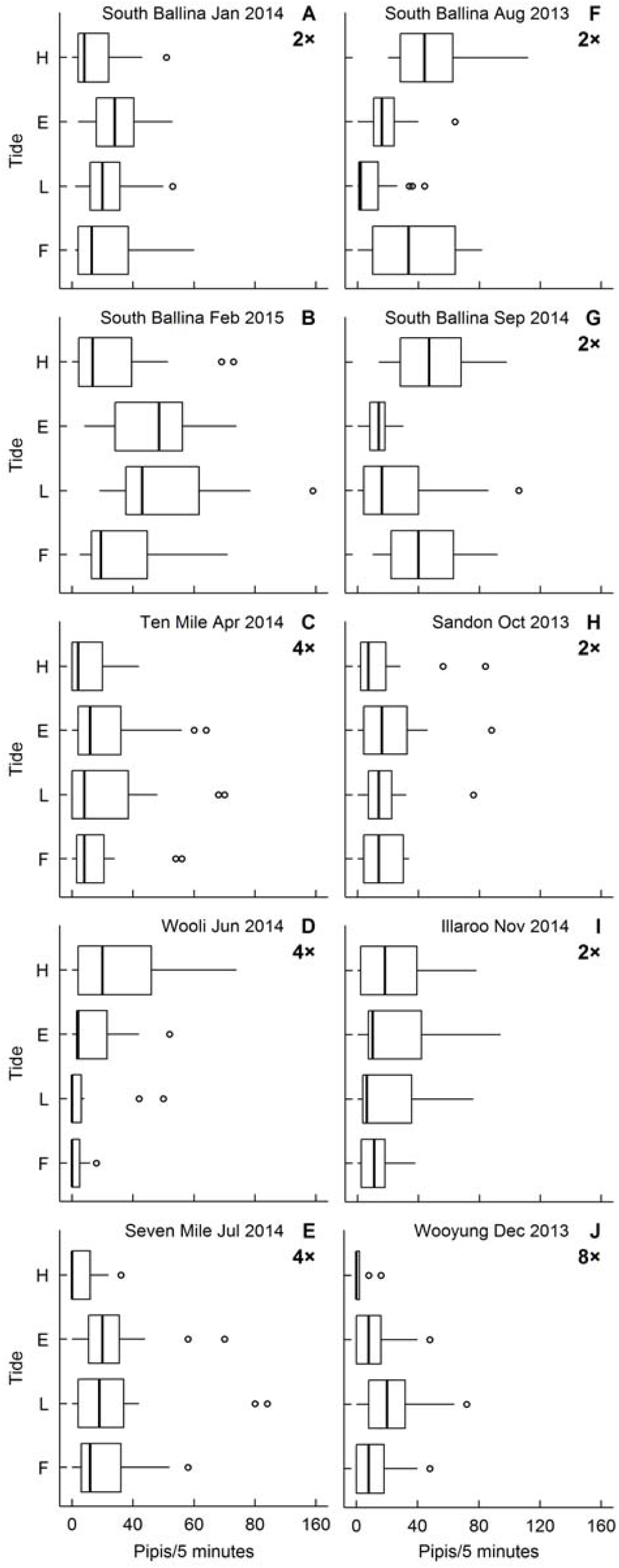
Box and whisker plots of feet digging clam counts by tidal stage (High, Ebb, Low, Flow). Counts for beaches with lower clam abundance are presented with expanded scales (2×, 4×, 8×). Plots are ordered by month. Two beaches were sampled in the months of January, April, August, December (Table 1) and the beach with the largest counts is presented. Boxes show the first quartile, median and third quartile, whiskers extend to a maximum of 1.5 times the interquartile range and data outside the whiskers are plotted as individual points.

The large number of feet digging plots sampled on each beach (mean 16 plots × 80 locations = 1280) delivered large length-frequency samples from this method (Figure 5). Length-frequency distributions from transect, clam band and feet digging samples were similar (*P*_*adj.*_ > 0.05) except for a recruitment event at South Ballina in September 2014, when feet digging missed most of the new recruits (*T*_*AD*_ = 50, *P* < 0.001; Fig 5f). After removing new recruits < 16 mm (Murray-Jones 1999) from those samples, no difference was detected (*T*_*AD*_ = 0.46, *P* = 0.24).

**Figure 5.**
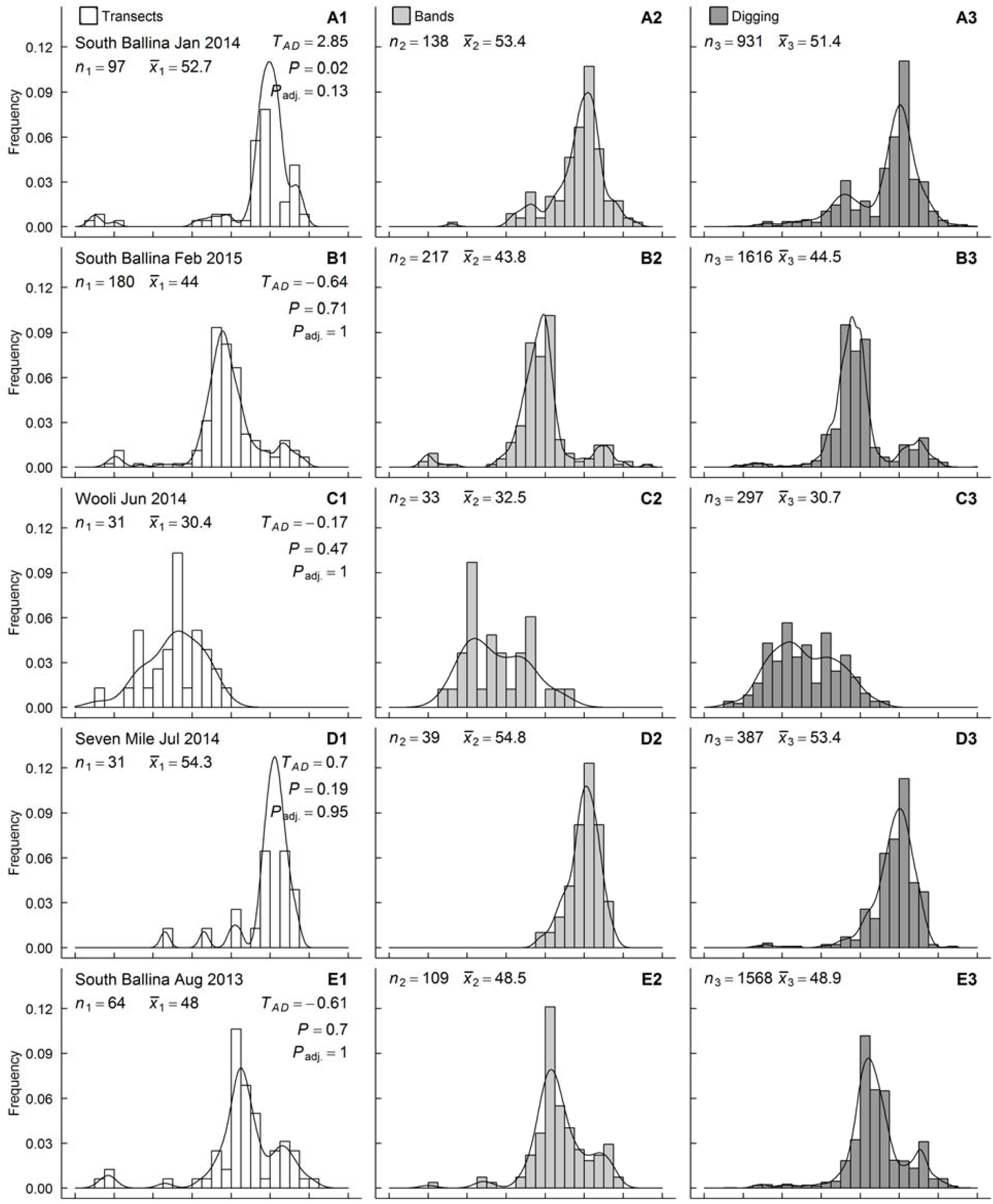
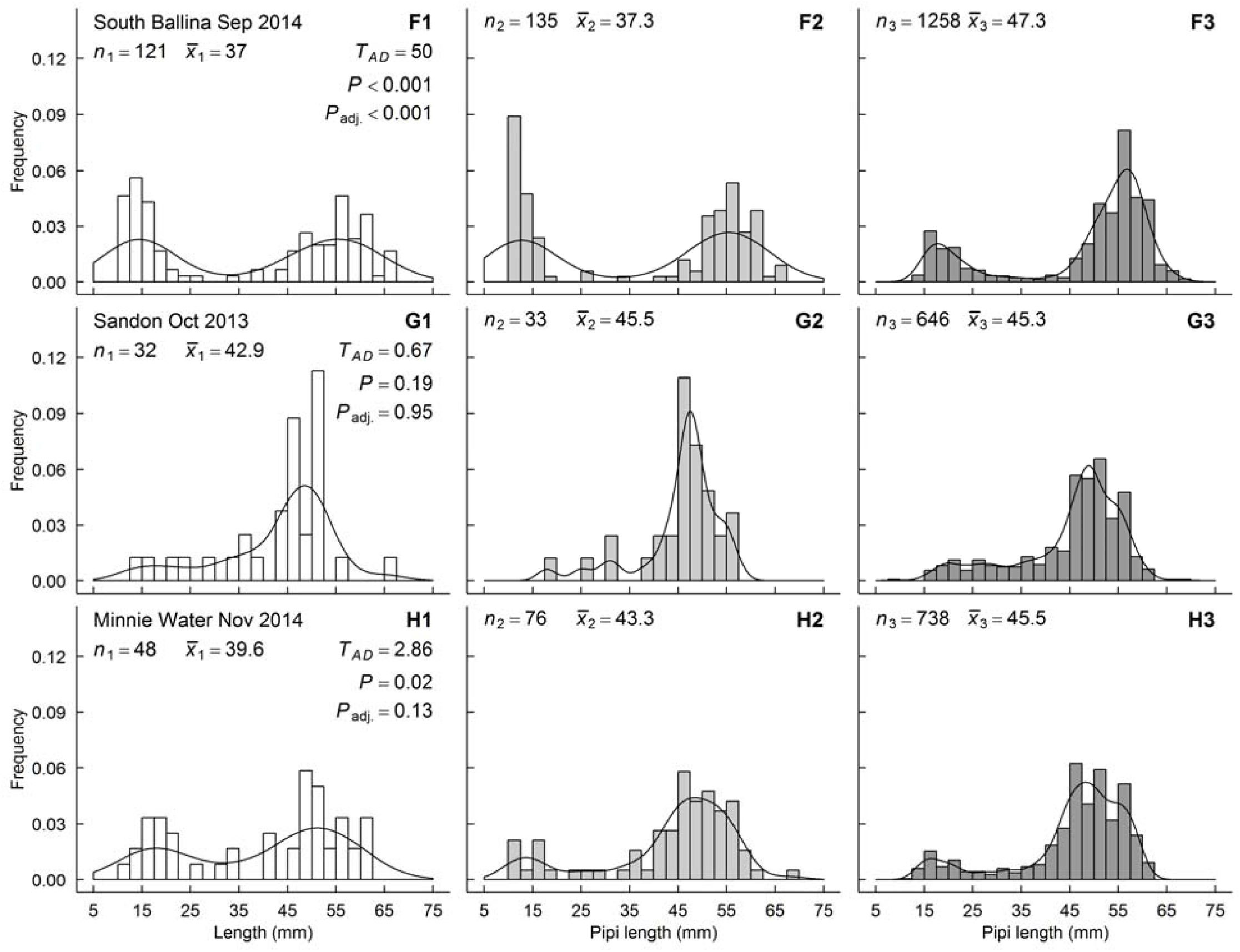
Length-frequency distributions for beaches with > 30 clams measured for each sampling method. Plots are grouped by beaches across rows and by methods (1 = transects; 2 = clam bands; 3 = feet digging) down columns. Beaches are ordered by month. Lines are kernel densities. Distributions were compared within beaches using *k*-sample Anderson-Darling tests (Scholz and Stephens 1987; standardised statistic *T*_*AD*_) with *P*-values adjusted for multiple comparisons (Holm 1979). Clams < 10 mm were not sampled by quadrats. A strong recruitment event is apparent in quadrat samples from South Ballina in September 2014 (f).

## 4. Discussion

The feet digging method was found to be satisfactory for measuring relative abundance and length-frequency distributions for *D. deltoides* ≥ 16 mm. As well, transect and clam band relative abundances correlated strongly with linear densities because the range of beach widths was narrow (Table 1).

Feet digging should be suitable for low cost relative abundance estimates, double sampling (*i.e.* feet digging counts can be “calibrated” with absolute abundances from transects) and busy beaches where human recreation (*e.g.* vehicle traffic) disturbs quadrat sampling. Feet digging was not effective for clams < 16 mm and is not suitable for studying recruitment. Feet digging is difficult in fine (< 0.25 mm), compact sand, although *D. deltoides* is usually rare on lower energy beaches with fine sand (pers. obs.).

It is recommended to distribute feet digging effort equally across the tidal cycle. Mean feet digging counts from any one of the four tidal stages (high, ebb, low and flow) correlated less strongly with linear densities compared to mean feet digging counts across all four stages. Totterman (2019b) reported seasonal low tide zonation patterns for *D. deltoides*, where clams typically aggregated in the intertidal zone in winter and in the low tide swash zone in summer. Seasonality in feet digging counts was less clear because tides are continuous, tide heights and wave heights are variable and clams in the swash zone are mobile.

Convenience sampling must be avoided. Swash zone sampling in Gray *et al.* (2014) and in Ferguson *et al.* (2015) was performed at low tide. Gray *et al.* (2016a) explained that low tide ±3 h was convenient for driving along beaches to access sampling locations. Totterman (2019b) raised concerns that Gray’s (2016b, 2016c) counts were biased by seasonal changes in low tide zonation of *D. deltoides*. Ferguson *et al.* (2015) stated in their methods that *D. deltoides* “are known to occur in a narrow band located 30–40 m below the mean high water mark within the swash zone” (on South Australian beaches). However, McLachlan *et al.* (1996) reported a broad, dense *D. deltoides* band from Goolwa Beach, extending across most of the intertidal zone and into the low tide swash zone. If only the low tide swash zone is to be sampled, it would be preferable that the bulk of the clam population is present in this zone and it must be assumed that the proportion of clams in the low tide swash zone is constant relative to the total number present across all zones.

Feet digging samples both across shore (between tides) and alongshore (between locations) variation in clam abundance. Accordingly, feet digging coefficients of variation averaged 1.2× larger and estimated sample sizes for constant relative standard error averaged 1.6× larger than those for quadrat-based methods. However, feet digging is faster (five minutes per location plus measurement time) than transect or clam band sampling (about two hours per location combined) and feet digging can continue throughout the tidal cycle.

In dense clam aggregations the observer can spend a large amount of time picking up (handling) clams relative to feet digging (searching) time. Like Holling’s (1959) “disc equation”, feet digging counts could approach an asymptote at very high densities, *i.e.* mean five minute counts would not be proportional to absolute abundance at very high densities. Effort could be better standardised by fixing the number of feet digging plots per sampling location (*e.g.* Gray *et. al.* 2014), keeping in mind that a reasonably large number of plots is recommended to cover the swash zone and average fine scale variation in clam abundance. Further studies could investigate the effects of number of locations sampled and number of feet digging plots per location on the precision of beach scale mean counts.

Gray *et al.* (2014) and Ferguson *et al.* (2015) did not compare their swash zone sampling relative abundances to quadrat-based results. Nonetheless, the assumption that relative abundances are proportional to true abundances may be more sensitive to the sampling design rather the sampling method. Key elements of a robust abundance sampling design for surf clams are: 1) identification of the target population, 2) definition of the sampling unit, 3) unbiased selection of sampling units and, 4) unbiased sampling within units.

Surf clam populations are naturally delimited by breaks in the beach habitat such as headlands, rivers and rocky intervals. Closely spaced quadrats or swash zone sampling plots are spatially autocorrelated and the correct sampling unit generally is the along shore sampling location (Millar and Anderson 2004). Beach scale sampling at replicate alongshore locations is necessary for mean abundance and accompanying variance estimates that refer to the clam population (Murray-Jones 1999). The Horvitz-Thompson estimator for the mean is:

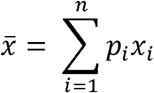

where *x*_*i*_ is the variable *x* measured on the *i*th sampling unit and *p*_*i*_ is the probability to sample the *i*th sampling unit (Albert *et al.* 2010). In a probabilistic sampling design, all *p*_*i*_ are equal and sum to one, which necessarily means that *p*_*i*_ = 1/*n*, where *n* is the sample size, giving the familiar arithmetic mean:

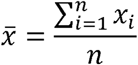

In a non-probabilistic design, some sampling units have no chance of being selected (*p*_*i*_ = 0; *e.g.* sampling only part of the beach) or their selection probability is variable and cannot be accurately determined (*e.g.* subjective selection of clam patches) so that estimates using the arithmetic mean can be biased. Unbiased sampling within sampling units (locations) is achieved by sampling the entire across shore distribution of clams (see above discussion of zonation and Totterman 2019b).

## Abbreviations

NSW: New South Wales;
CPUE: catch per unit effort

## Acknowledgements

Catch, measure and release sampling of *D. deltoides* was permitted under Section 37 of the Fisheries Management Act 1994, permit number P13/0024-1.0 issued by Fisheries NSW, Department of Primary Industries, NSW Government. Hadley Wickham is thanked for the ggplot2 graphics package for R (Wickham 2009).

